# The N-terminal Tail of *C. elegans* CENP-A Interacts with KNL-2 and is Essential for Centromeric Chromatin Assembly

**DOI:** 10.1101/2020.12.28.424576

**Authors:** Christian de Groot, Jack Houston, Bethany Davis, Adina Gerson-Gurwitz, Joost Monen, Karen Oegema, Andrew K. Shiau, Arshad Desai

**Author notes:** co-first authors (Christian de Groot, Jack Houston). Corresponding author, Address: CMM-E Rm 3052, 9500 Gilman Dr, La Jolla, CA 92093-0653.

## Abstract

Centromeres are epigenetically defined by the presence of the centromere-specific histone H3 variant CENP-A. A specialized loading machinery, including the histone chaperone HJURP/Scm3, participates in CENP-A nucleosome assembly. However, Scm3/HJURP is missing from multiple lineages, including nematodes, which rely on a CENP-A-dependent centromere. Here, we show that the extended N-terminal tail of *C. elegans* CENP-A contains a predicted structured region that is essential for centromeric chromatin assembly. Removal of this region of the CENP-A N-Tail prevents loading, resulting in failure of kinetochore assembly and defective chromosome condensation. By contrast, the N-Tail mutant CENP-A localizes normally in the presence of endogenous CENP-A. The portion of the N-Tail containing the predicted structured region binds to KNL-2, a conserved SANTA and Myb domain-containing protein (referred to as M18BP1 in vertebrates), that is specifically involved in CENP-A chromatin assembly. This direct interaction is conserved in the related nematode *C. briggsae,* despite divergence of the N-Tail and KNL-2 primary sequences. Thus, the extended N-Tail of CENP-A is essential for CENP-A chromatin assembly in *C. elegans* and partially substitutes for the function of Scm3/HJURP, in that it mediates an interaction of the specialized histone fold of CENP-A with KNL-2. These results highlight an evolutionary variation on centromeric chromatin assembly in the absence of a dedicated CENP-A-specific chaperone/targeting factor of the Scm3/HJURP family.

## INTRODUCTION

Centromeres are specialized chromosomal loci that direct chromosome segregation. In most species, active centromeres are defined by the presence of CENP-A, a histone variant that replaces histone H3 in centromeric nucleosomes (Kixmoeller et al., 2020; Mitra et al., 2020). CENP-A provides the physical foundation for assembly of the kinetochore, a multiprotein complex mediating spindle microtubule attachment to chromosomes (Musacchio and Desai, 2017). The cues leading to the centromere-restricted localization of CENP-A are being actively investigated. The underlying centromeric DNA is not conserved and, with the exception of budding yeasts, neither necessary nor sufficient to propagate CENP-A chromatin (Allshire and Karpen, 2008; McKinley and Cheeseman, 2016).

A segment of the histone fold domain (HFD) of CENP-A, known as the CENP-A targeting domain (CATD), when transferred into canonical histone H3 is sufficient to confer centromere localization (Black et al., 2004). The CATD of CENP-A interacts with a CENP-A-specific histone chaperone, known as <u>H</u>olliday <u>j</u>unction <u>r</u>epair <u>p</u>rotein (HJURP) in vertebrates and Scm3 in fungi (Dunleavy et al., 2009; Foltz et al., 2009). This interaction is essential for CENP-A centromere targeting during mitotic exit (Foltz et al., 2009; Jansen et al., 2007). However, HJURP/Scm3 is not conserved in all species that build centromeres using CENP-A, including insects and nematodes (McKinley and Cheeseman, 2016). In *Drosophila melanogaster,* there is compelling evidence that the unrelated protein Cal1 acts as a functional homologue of HJURP/Scm3 (Chen et al., 2014; Erhardt et al., 2008; Mellone et al., 2011). By contrast, in *C. elegans,* no HJURP/Scm3-like activity has been identified to date.

In addition to promoting CENP-A nucleosome assembly, HJURP/Scm3 proteins target the CENP-A/H4-HJURP/Scm3 prenucleosomal complex to the specific location of the centromere by interaction with centromeric DNA/chromatin-bound targeting factors. In budding yeast, the CBF3 complex specifically recognizes centromeric DNA and its subunit Ndc10 interacts with HJURP/Scm3 of the prenucleosomal complex to localize new CENP-A nucleosome assembly (Cho and Harrison, 2011). Outside of budding yeasts, where the CBF3 complex is not present, the Mis18 complex is the primary candidate for recognizing existing centromeric chromatin domains and targeting the deposition of new CENP-A via an interaction with HJURP/Scm3. The Mis18 complex is composed of Mis18 and/or KNL-2/M18BP1, depending on the species: Mis18α, Mis18β & M18BP1 in humans (Fujita et al., 2007); Mis18 only in *S. pombe* (Hayashi et al., 2004; Pidoux et al., 2009; Williams et al., 2009); KNL-2 only in *C. elegans* (Maddox et al., 2007) and *Arabidopsis* (Lermontova et al., 2013). *S. pombe* Mis18 and human Mis18α/β interact with HJURP/Scm3 *in vitro* (Pan et al., 2019; Pidoux et al., 2009; Wang et al., 2014) and Mis18 complex-mediated CENP-A recruitment can be bypassed by artificial tethering of HJURP/Scm3 to chromatin (Barnhart et al., 2011; Foltz et al., 2009; Ohzeki et al., 2012). In human cells, centromere localization of the Mis18 complex precedes that of the CENP-A-H4-HJURP prenucleosomal complex during mitotic exit-coupled new CENP-A chromatin assembly (Foltz et al., 2009; Jansen et al., 2007). The Mis18 complex is proposed to recognize existing centromeric chromatin at least in part by binding to CENP-C, the reader of CENP-A nucleosomes that directs kinetochore assembly (Kato et al., 2013; Moree et al., 2011). However, in *C. elegans,* KNL-2 localizes to chromatin independently of CENP-C (Maddox et al., 2007) and, even in non-mammalian vertebrates, the Mis18 complex directly recognizes CENP-A nucleosomes (French et al., 2017; Hori et al., 2017). KNL-2/M18BP1 family proteins contain a conserved Myb-like DNA binding domain and a SANTA domain whose functions independent of CENP-C association are unclear (French and Straight, 2019; Maddox et al., 2007; Ohzeki et al., 2012; Zhang et al., 2006). Interestingly, in fungi where KNL-2/M18BP1 proteins are absent, Myb domains can be found in HJURP/Scm3 proteins (Sanchez-Pulido et al., 2009), suggesting potential fusion of multiple functions within a single polypeptide.

Here, we investigate CENP-A chromatin in *C. elegans,* which requires KNL-2/M18BP1 for its assembly but lacks an HJURP/Scm3 family member. *C. elegans* is holocentric, with condensed mitotic chromosomes having two “stripes” of CENP-A chromatin—one per sister chromatid—on geometrically opposite surfaces (Maddox et al., 2007; Melters et al., 2012). Genomic approaches have localized CENP-A and KNL-2 to broad permissive domains in the *C. elegans* genome that are transcriptionally inactive (Gassmann et al., 2012), although restriction to specific sites has also been suggested (Steiner and Henikoff, 2014). We focused on the unusually long amino-terminal tail (N-Tail) of *C. elegans* CENP-A, which, unlike the N-termini of CENP-A from other species, is predicted to harbor a region with α-helical secondary structure. Analysis of the divergent N-Tails of CENP-A family members have implicated them in kinetochore assembly and epigenetic stability of centromeric chromatin (Fachinetti et al., 2013; Folco et al., 2015; Ravi et al., 2010). Employing single copy, targeted transgene insertion to replace endogenous CENP-A, we find that the N-Tail of *C. elegans* CENP-A is essential for CENP-A loading, and we link this essential function to a direct interaction between the N-Tail and the loading factor KNL-2/M18BP1. Interaction of the extended N-Tail of *C. elegans* CENP-A to KNL-2/M18BP1 represents an evolutionary variation to HJURP/Scm3-mediated targeting of the specialized histone fold of CENP-A to centromeres.

## RESULTS

### The *C. elegans* CENP-A N-Tail has a Predicted Structured Region that is Essential for Viability

The *C. elegans* CENP-A N-Tail is unusually long at 189 amino acids and, based on computational analysis (performed using PSIPRED; http://bioinf.cs.ucl.ac.uk/psipred/), is predicted to be α-helical in the first 100 amino acids and unstructured afterwards (**Fig. 1A**). The presence of a structured region in the N-tail is unexpected as the CENP-A tail is often short (e.g. in fission yeast or humans, where it is 20 and 39 aa, respectively) and, even in other species with extended CENP-A N-Tails—such as *D. melanogaster* (123 aa) or *S. cerevisiae* (130 aa)—are not predicted to have any secondary structure (**Fig. 1A**).

**Figure 1.**
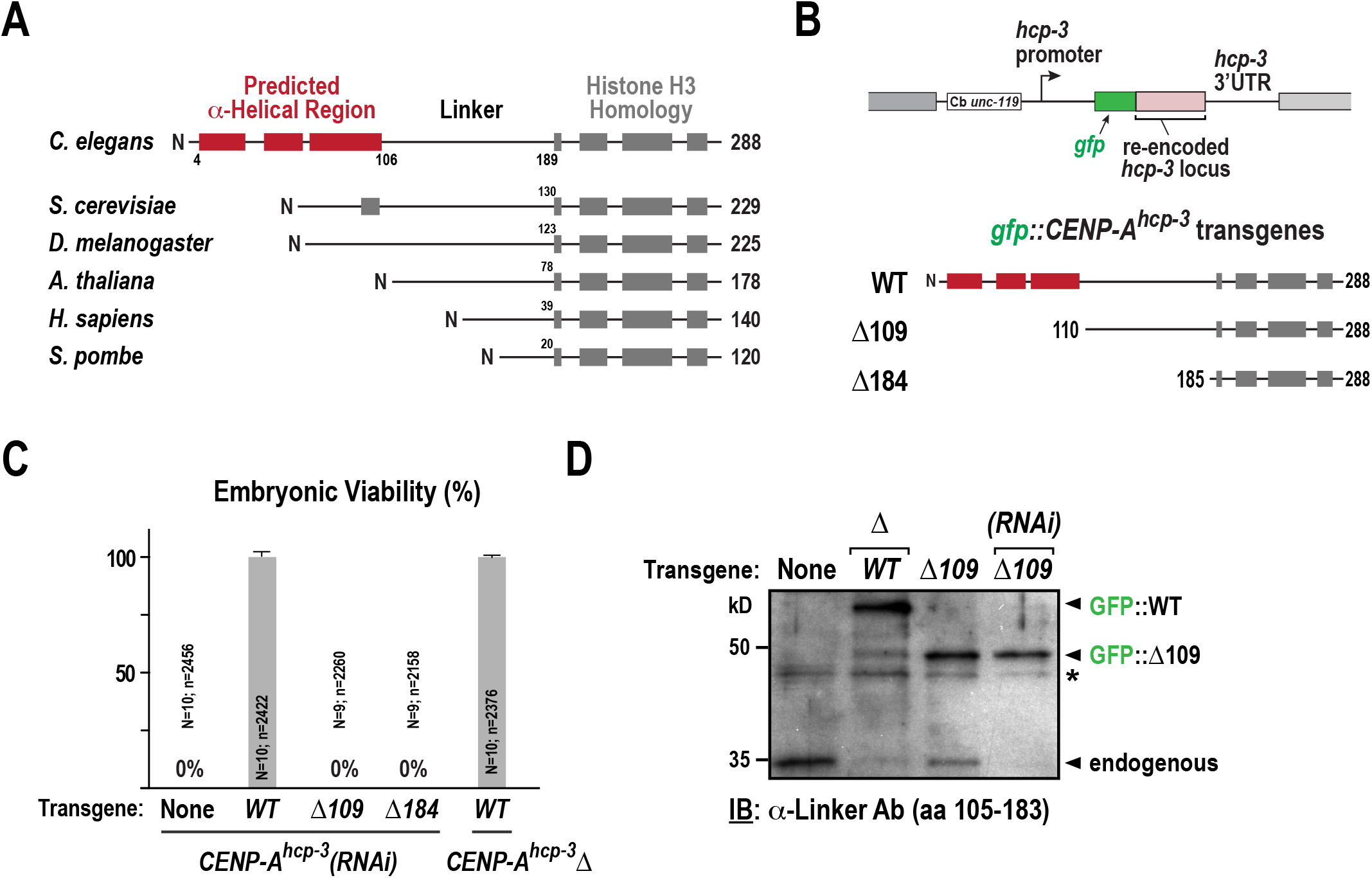
The extended N-Tail of *C. elegans* CENP-A^HCP-3^ contains a predicted structured region that is essential for viability. **(A)** Secondary structure predictions of CENP-A from different model organism species. Secondary structure predictions were generated using PsiPred. Predicted alpha helical segments are indicated as boxes. The histone fold domain (HFD) is marked in grey. **(B)** Schematic of RNAi-resistant *gfp::CENP-A^hcp-3^* single copy transgene insertions on Chromosome II. The three variants of CENP-A^HCP-3^ expressed from single copy transgene insertions are indicated below. **(C)** Embryo viability analysis for the indicated conditions. *N* refers to the number of worms and *n* to the total number of embryos scored. Error bars are the SEM. **(D)** Anti-CENP-A^HCP-3^ immunoblot performed using an antibody raised to the linker region (aa105-183) showing expression levels of WT GFP-CENP-A^HCP-3^ and the Δ109 N-Tail truncation mutant in the presence and absence of endogenous CENP-A^HCP-3^ (Δ indicates homozygous *CENP-A^hcp-3^* deletion mutant; *(RNAi)* indicates *CENP-A^hcp-3^(RNAi)).* Asterisk (*) marks a background band that serves as a loading control.

To test the functional significance of the predicted structured region of the *C. elegans* CENP-A N-Tail, we developed a transgene-based system to replace endogenous CENP-A (named HCP-3 and referred to here as CENP-A^HCP-3^) in *C. elegans* with engineered N-Tail mutants. In brief, the nucleotide sequence of the *CENP-A^hcp-3^* coding region was altered to maintain the native amino acid sequence while enabling selective RNAi-mediated depletion of endogenous CENP-A^HCP-3^; in addition, an N-terminal GFP tag was added to monitor localization (**Fig. S1**). The wildtype, as well as two mutant transgenes (Δ109, which removes the predicted structured region, and Δ184, which removes the majority of the N-Tail), were inserted in single copy at a fixed genomic location harboring a Mos transposon insertion (**Fig. 1B; Fig. S1**). The re-encoded *gfp::CENP-A^hcp-3^* transgene fully rescued embryonic lethality observed following depletion of endogenous CENP-A^HCP-3^ by RNAi as well as the lethality of a deletion mutant *(hcp-3(ok1892),* referred to as *CENP-A^hcp-3^Δ;* **Fig 1C**). By contrast, deletion of the predicted structured region (Δ109) and of the majority of the N-Tail (Δ184) resulted in fully penetrant embryonic lethality (**Fig. 1C**). Immunoblotting with an antibody raised to the unstructured linker (amino acids 105-183 of the N-Tail) indicated that the Δ109 mutant was expressed similarly to endogenous CENP-A^HCP-3^ and the transgene-encoded WT GFP::CENP-A^HCP-3^ (**Fig. 1D**). Hence, the observed lethality is not because the Δ109 N-Tail mutant is not expressed. We therefore conclude that the predicted structured region of the N-Tail of CENP-A is essential for viability of *C. elegans* embryos.

### The CENP-A^HCP-3^ N-Tail Deletion Mutant Exhibits a Kinetochore-Null Phenotype in OneCell Embryos

We next assessed the phenotype observed when endogenous CENP-A^HCP-3^ was replaced by the Δ109 N-Tail mutant. We crossed an mCherry::H2b marker into strains harboring singlecopy transgenes expressing WT or Δ109 GFP::CENP-A^HCP-3^, depleted endogenous CENP-A^HCP-3^ by RNAi, and imaged one-cell embryos. As a control, we also depleted CENP-A^HCP-3^ in the absence of any transgene. Depletion of CENP-A resulted in the characteristic kinetochore-null phenotype, with two clusters of chromatin—one from each pronucleus— instead of a metaphase plate, and a failure of segregation (**Fig. 2A**; (Desai et al., 2003; Oegema et al., 2001). This severe phenotype was fully rescued by transgene-encoded RNAi-resistant WT GFP::CENP-A^HCP-3^. By contrast, the observed phenotype for the Δ109 N-Tail mutant was similar to that of removal of CENP-A^HCP-3^ (**Fig. 2A**). Thus, deletion of the first 109 amino acids of the N-Tail of CENP-A^HCP-3^ results in a chromosome segregation phenotype that is equivalent to CENP-A^HCP-3^ removal in the *C. elegans* embryo.

**Figure 2.**
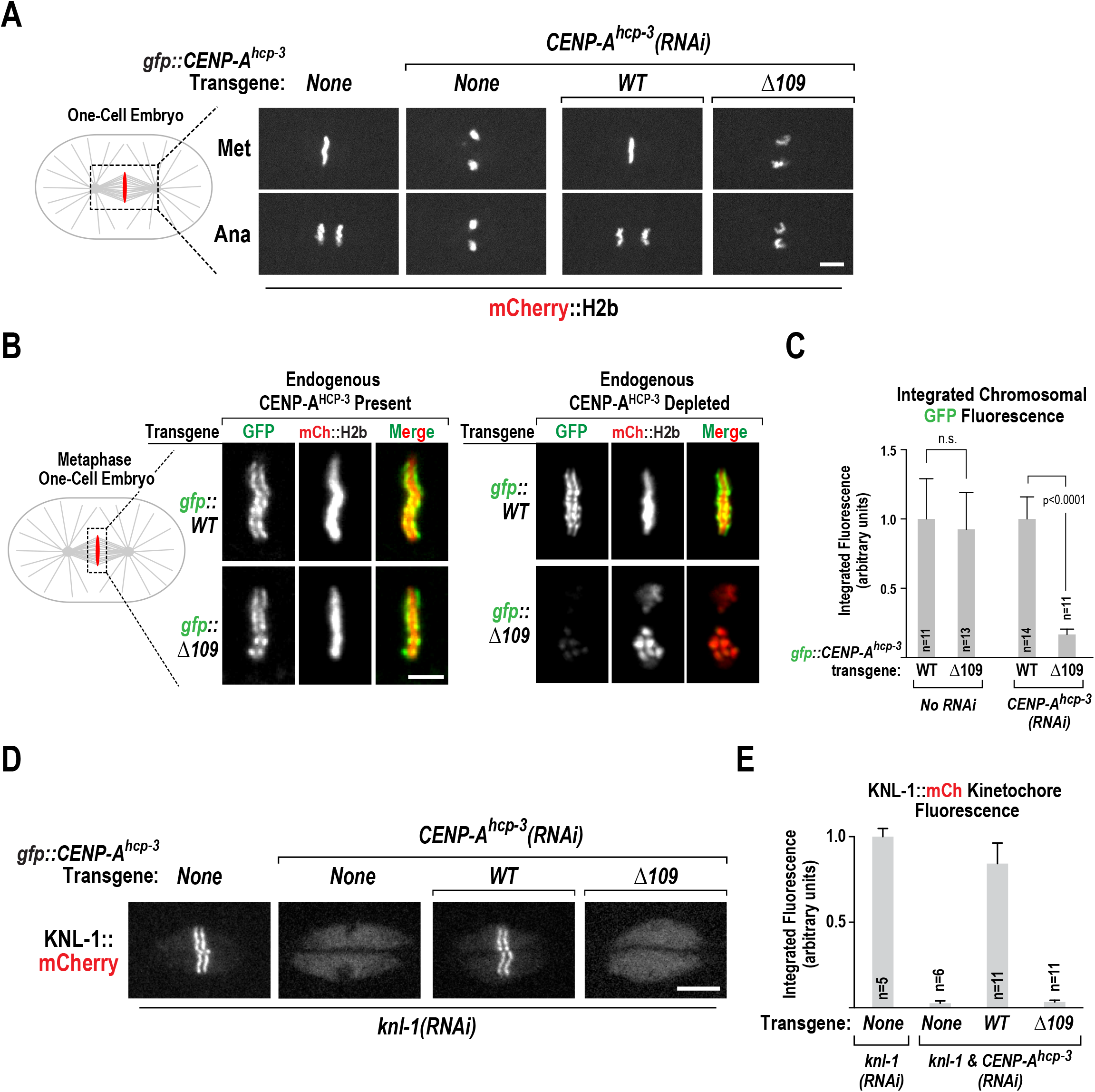
Deletion of the predicted α-helical region of the CENP-A^HCP-3^ N-Tail results in a kinetochore-null phenotype and failure to accumulate on mitotic chromatin. **(A)** mCherry::H2b images from timelapse sequences for the indicated conditions in metaphase and anaphase-stage one-cell embryos. Similar results were obtained in at least 10 embryos per condition. Scale bar, 5 μm. **(B)** Images of WT and Δ109 GFP::CENP-^HCP-3^ in metaphase stage embryos expressing mCherry-H2b in the presence (*left set of panels*) or absence (*right set of panels*) of endogenous CENP-A^HCP-3^. Scale bar, 2.5 μm. **(C)** Quantification of integrated chromosomal GFP intensity in metaphase stage embryos for the indicated conditions. t-tests were used to assess if indicated pair-wise comparisons were significantly different. Error bars are the SD. **(D)** Images of KNL-1::mCherry, expressed from an integrated single copy RNAi-resistant transgene, in metaphase stage one-cell embryos for the indicated conditions; note that endogenous KNL-1 was depleted in all cases. Scale bar, 5 μm. **(E)** Quantification of integrated KNL-1::mCherry kinetochore intensity in metaphase stage embryos for the indicated conditions. Error bars are the SD.

### The CENP-A^HCP-3^ N-Tail Deletion Mutant Does Not Accumulate on Chromatin and Fails to Support Kinetochore Assembly

Stable incorporation of CENP-A into chromatin in yeast and humans involves a region of the histone fold, referred to as the CATD, which is specifically bound by the chaperone Scm3/HJURP (Black et al., 2007; Cho and Harrison, 2011; Hu et al., 2011). In these species, the N-tail is not essential for CENP-A centromere targeting, and alterations of the N-Tail do not phenocopy loss of CENP-A (Chen et al., 2000; Fachinetti et al., 2013; Folco et al., 2015). The similar phenotypes observed for CENP-A^HCP-3^ removal and for the Δ109 N-Tail mutant of *C. elegans* CENP-A suggested that the Δ109 N-Tail mutant, in contrast to the N-Tail mutants in other species, does not accumulate on centromeric chromatin. To test this idea, we imaged WT and Δ109 GFP::CENP-A^HCP-3^ in a strain co-expressing mCherry::H2b, and quantified the GFP signal on metaphase chromosomes. In the presence of endogenous CENP-A^HCP-3^, both WT and Δ109 GFP::CENP-A^HCP-3^ localized to the diffuse kinetochores on the poleward faces of the holocentric mitotic chromosomes (**Fig. 2B**) and quantification of fluorescence intensity indicated equivalent localization of both (**Fig. 2C**). However, in the absence of endogenous CENP-A^HCP-3^, localization of Δ109 GFP::CENP-A^HCP-3^ was greatly reduced relative to WT GFP::CENP-A^HCP-3^ (**Fig. 2B,C**). Thus, Δ109 GFP::CENP-A^HCP-3^ fails to localize to chromatin on its own.

The absence of the Δ109 CENP-A^HCP-3^ mutant in chromatin should result in a kinetochore assembly defect. To confirm that this was indeed the case, we analyzed the localization of an RNAi-resistant mCherry-fusion of KNL-1, an outer kinetochore scaffold protein (Desai et al., 2003; Espeut et al., 2012). We introduced the transgene expressing this fusion into the strain expressing either WT or Δ109 GFP::CENP-A^HCP-3^, depleted endogenous KNL-1 and CENP-A^HCP-3^, and imaged and quantified the KNL-1::mCherry signal. This analysis revealed loss of KNL-1::mCherry localization in the Δ109 GFP::CENP-A^HCP-3^ mutant was analogous to the CENP-A^HCP-3^ depletion (**Fig. 2D,E**). Thus, the Δ109 CENP-A^HCP-3^ mutant does not form centromeric chromatin, resulting in a failure in kinetochore assembly.

### The Failure of the CENP-A^HCP-3^ N-Tail Deletion Mutant to Accumulate on Chromatin is Not Due to a Kinetochore Assembly Defect

CENP-A^HCP-3^ is essential for kinetochore assembly (Oegema et al., 2001). The severe reduction of Δ109 CENP-A^HCP-3^ on condensed chromatin could be either due to a defect in its loading or a secondary consequence of its inability to support kinetochore assembly. To distinguish between these possibilities, we prevented kinetochore assembly by depleting CENP-C^HCP-4^ or KNL-1, which are recruited downstream of CENP-A^HCP-3^ to build the outer kinetochore (Desai et al., 2003; Oegema et al., 2001), and analyzed the effect on WT GFP::CENP-A^HCP-3^ chromatin accumulation; endogenous CENP-A^HCP-3^ was also depleted. As expected from prior work, both CENP-C^HCP-3^ and KNL-1 depletions resulted in a kinetochore null phenotype. However, CENP-A^HCP-3^ accumulation on chromatin was similar to WT controls (**Fig, 3A,B**), indicating that a failure in kinetochore assembly is not the reason for the loss of CENP-A^HCP-3^ chromatin localization. Thus, the absence of Δ109 CENP-A^HCP-3^ on chromatin is likely due to a defect in its loading, rather than a consequence of its inability to support kinetochore assembly.

**Figure 3:**
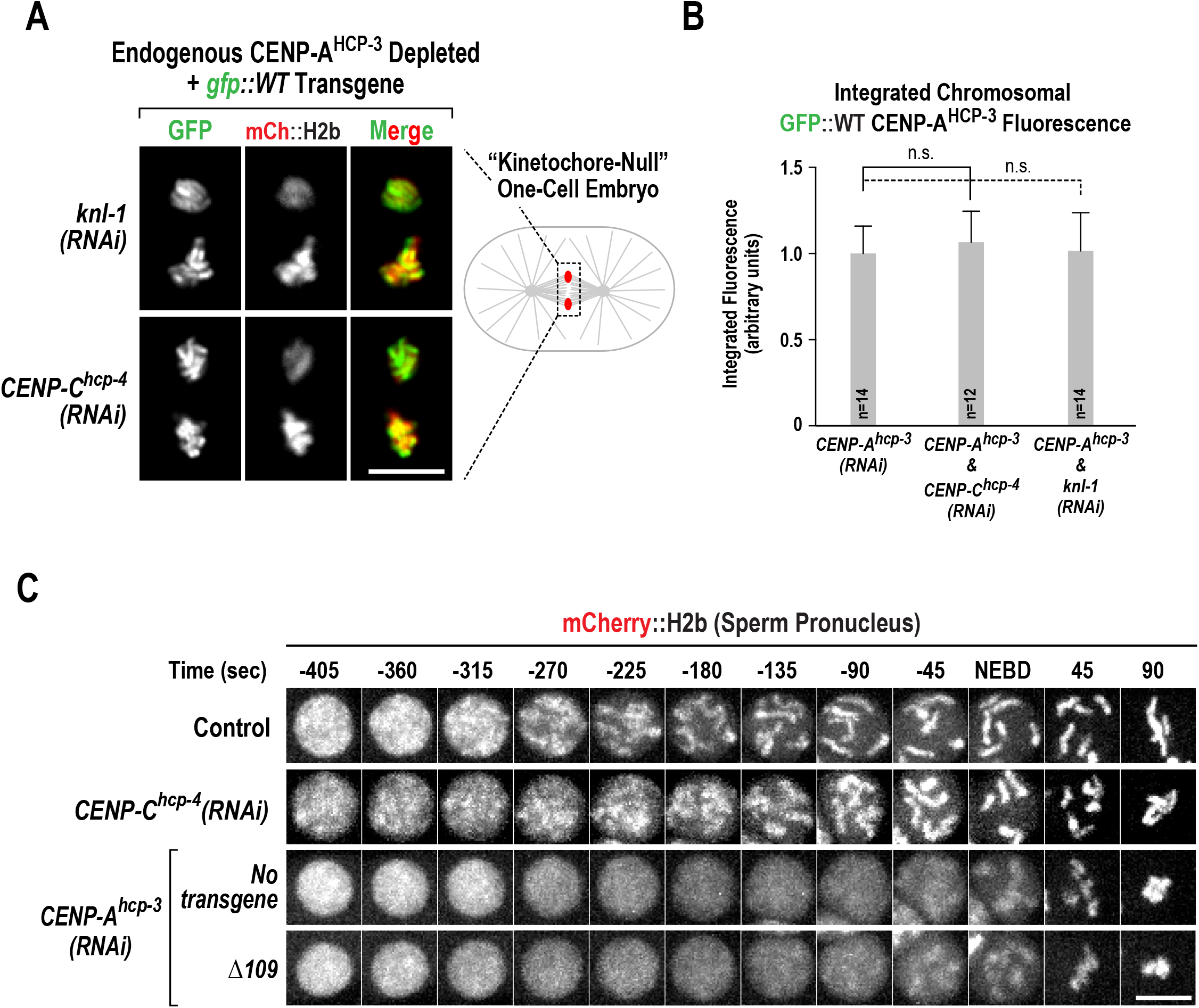
Inability of Δ109 CENP-A^HCP-3^ to accumulate on chromatin is not due to failure of kinetochore assembly. **(A)** Images of WT GFP::CENP-A^HCP-3^ in metaphase stage embryos also expressing mCherry::H2b that were depleted of endogenous CENP-A^HCP-3^ and KNL-1 (*top*) or CENP-C^HCP-4^ *(bottom).* Scale bar, 5 μm. **(B)** Quantification of integrated chromosomal WT GFP::CENP-A^HCP-3^ intensity in metaphase stage embryos for the indicated conditions. The *CENP-A^hcp-3^(RNAi)* alone value is the same as in *Fig. 2C.* Error bars are the SD. t-tests were employed to assess statistical significance of indicated pair-wise comparisons. **(C)** Images of mCherry::H2b in sperm pronuclei from timelapse sequences for the indicated conditions. Times are in seconds after nuclear envelope breakdown (NEBD). Similar results were observed in at least 10 embryos filmed per condition. Scale bar, 5 μm.

CENP-A^HCP-3^ depletion, in addition to the kinetochore-null phenotype, also leads to defects in condensation of the holocentric *C. elegans* chromosomes (Maddox et al., 2007; Maddox et al., 2006). By contrast, preventing kinetochore formation by depletion of CENP-C^HCP-4^ does not result in a severe condensation defect (Maddox et al., 2006). We therefore compared chromosome condensation in CENP-A^HCP-3^ and CENP-C^HCP-4^ depletion to that in the Δ109 CENP-A^HCP-3^ tail mutant. This analysis focused on sperm pronuclei as they are formed prior to injection of the dsRNA employed to deplete endogenous CENP-A^HCP-3^ and are therefore free of potential meiotic defects (Maddox et al., 2006). Chromosome condensation in the Δ109 CENP-A^HCP-3^ mutant resembled that resulting from CENP-A^HCP-3^ depletion and not CENP-C^HCP-4^ depletion, providing additional support that the Δ109 mutant is compromised for its loading onto chromatin. Taken together, the results from these two distinct assays argue that the severe reduction of Δ109 mutant of CENP-A^HCP-3^ on chromatin is not due to an inability to support kinetochore assembly but due to a defect in its loading.

### The Predicted α-Helical Region of the CENP-A Tail Interacts with the CENP-A Loading Factor KNL-2

The above results implicate the predicted structured region of the N-Tail of *C. elegans* CENP-A^HCP-3^ in its loading on chromatin. In species with Scm3/HJURP, a key step in the loading reaction is the interaction of the Scm3/HJURP-CENP-A complex with the Mis18 complex, which includes the Myb domain-containing protein KNL-2 (also known as Mis18BP1, based on its association with Mis18α/β in human cells (French et al., 2017; Fujita et al., 2007; Maddox et al., 2007; Pan et al., 2019; Wang et al., 2014); to date, a Mis18α/β homolog has not been identified in *C. elegans*). In addition, Scm3/HJURP functions as a chaperone for assembly of CENP-A nucleosomes (Dunleavy et al., 2009; Foltz et al., 2009; Pidoux et al., 2009; Shuaib et al., 2010; Williams et al., 2009).

To determine how the N-Tail of CENP-A^HCP-3^ contributes to its loading, we tested if it interacts with KNL-2. KNL-2 family proteins are characterized by a conserved Myb-like DNA binding domain and a predicted folded N-terminal domain referred to as the SANTA domain (**Fig. 4A**; (Maddox et al., 2007; Zhang et al., 2006); in addition, they possess an acidic/aromatic tail at the C-terminus (**Fig. S2A**). Using yeast two-hybrid analysis, we observed a robust interaction between a segment of the middle region of KNL-2 (residues 267 to 470; predicted to be unstructured) and the predicted α-helical region of the N-Tail of CENP-A^HCP-3^; an interaction between KNL-2 and CENP-A^HCP-3^ was also reported in a large-scale two-hybrid screen of proteins essential for *C. elegans* embryogenesis (Boxem et al., 2008). This interaction was not observed with full-length KNL-2 but, as no interaction has been observed with this fusion, this may be a false negative due to the full-length protein not being properly expressed/folded in yeast. Importantly, an interaction between similar regions of CENP-A^HCP-3^ and KNL-2 was also observed for the *C. briggsae* proteins (**Fig. 4B**), despite primary sequence divergence (23.7% identity/43.5 % similarity for CENP-A^HCP-3^ N-Tail & 42.9% identity/55.1% similarity for KNL-2 middle region; **Fig. S2B**). The interaction was species-specific, as the *C. briggsae* CENP-A^HCP-3^ N-Tail did not interact with *C. elegans* KNL-2 middle region and vice versa (**Fig. 4B**).

**Figure 4.**
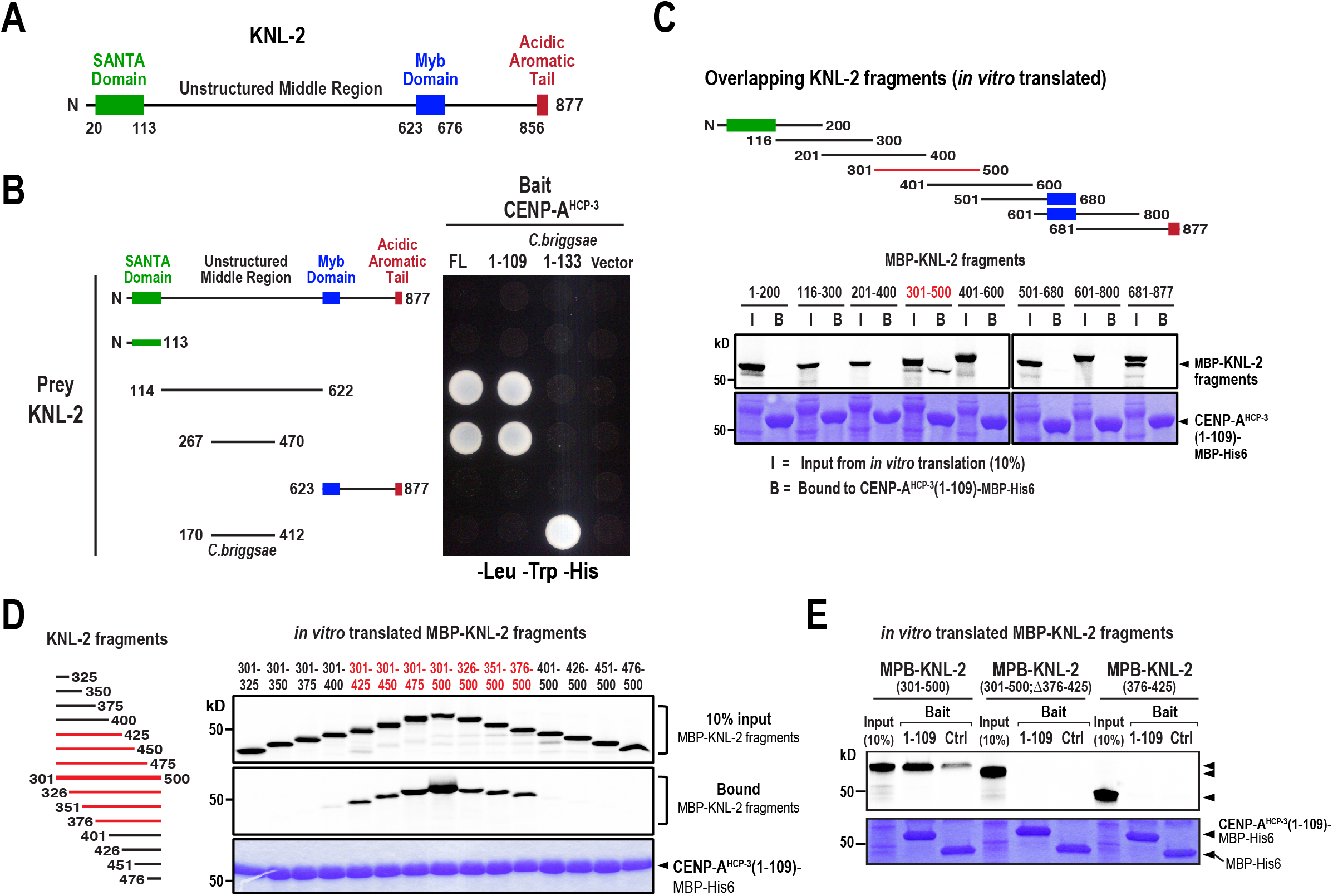
The N-Tail of CENP-A^HCP-3^ interacts with an unstructured middle region of KNL-2. **(A)** Domain structure of KNL-2. The presence of the SANTA domain (PF09133) and the Myb domain (also known as the SANT domain; PF00249) is conserved among the KNL-2/M18BP1 protein family. No secondary structure elements are predicted in the middle region of KNL-2. **(B)** Yeast two-hybrid analysis of CENP-A^HCP-3^ N-Tail and KNL-2. The bait CENP-A^HCP-3^ N-Tail fusions are listed on top and the prey KNL-2 fusions are listed on the left. **(C)-(E)** Biochemical analysis of the CENP-A^HCP-3^-KNL-2 interaction. Nickel-immobilized recombinant CENP-A^HCP-3^(1-109)-MBP-His6 was used to pull-down indicated reticulocyte lysate-expressed S^35^-labeled MBP-KNL-2 fragments. In (*C*), S^35^-autoradiogram (*top*) shows Input (I) and bead-bound (B) KNL-2 fragments; Coomassie staining (*bottom*) shows input lysate and CENP-A^HCP-3^(1-109)-MBP-His6 bait. In (*D*), fragments of the 301-500 amino acid region of KNL-2 tested for binding to CENP-A^HCP-3^(1-109) are schematized on the left. S^35^- autoradiogram (*top*) shows reticulocyte lysate-expressed MBP-KNL-2 fragments; S^35^- autoradiogram *(middle)* shows bound KNL-2 fragments; Coomassie staining *(bottom)* shows CENP-A^HCP-3^ N-Tail bait. In (*E*), indicated KNL-2 fragments were tested for binding to control (MBP-His6) and CENP-A^HCP-3^(1-109)-MBP-His6 baits. S^35^-autoradiogram (*top*) shows input and bound fragments; Coomassie staining (*bottom*) shows input lysates and baits.

To confirm the two-hybrid interaction, we performed pull-down assays with a purified MBP-His6 fusion of the CENP-A^HCP-3^ N-Tail immobilized on nickel agarose and *in vitro-* translated MBP-KNL-2 fragments. We first screened a series of overlapping fragments of KNL-2 and found that, consistent with the yeast two-hybrid results, a fragment containing residues 301-500 interacted with the N-tail (**Fig. 4C,E**). We next analyzed a series of truncated fragments in this region and found that a central region of 50 amino acids (376-425) was essential for the interaction (**Fig. 4D,E**). However, this 50 amino acid region on its own did not interact with the CENP-A^HCP-3^ N-Tail (**Fig. 4E**), suggesting that residues on either side are important for the observed interaction. We conclude that the predicted α-helical region of the N-Tail of CENP-A^HCP-3^ that is important for chromatin loading *in vivo* interacts directly with the CENP-A^HCP-3^ loading factor KNL-2 *in vitro* and that this interaction is conserved in a related nematode species with a significantly diverged N-Tail sequence.

## DISCUSSION

Here, we investigated how CENP-A chromatin is assembled in the absence of a HJURP/Scm3 family protein in *C. elegans*. We found that the unusually long N-tail of *C. elegans* CENP-A, which contains a predicted α-helical region, is required for CENP-A loading onto chromatin. By contrast, the divergent N-Tails of CENP-A family members have been proposed to contribute to kinetochore assembly and to epigenetic stability of centromeric chromatin (Fachinetti et al., 2013; Folco et al., 2015; Ravi et al., 2010), but have not been implicated in assembly of CENP-A chromatin. By comparing deletion of the predicted structured region of the CENP-A N-Tail to two other perturbations that prevent kinetochore assembly, we show that the absence of N-Tail-mutant CENP-A^HCP-3^ on chromatin is due to a failure in loading and not a consequence of defective kinetochore assembly. In addition, as the N-Tail-mutant CENP-A^HCP-3^ localizes normally in the presence of endogenous CENP-A^HCP-3^, the absence of localization cannot be attributed to misfolding or inability to interact with histone H4. Thus, the N-Tail effectively acts as an intramolecular-targeting signal, analogous to the CATD in HJURP/Scm3 containing species.

HJURP/Scm3 has two roles—one is to act as a chaperone promoting assembly of CENP-A nucleosomes, and the second is to target this assembly reaction to centromeric chromatin through an interaction with centromere recognition factors (Ndc10 in budding yeast, Mis18 in fission yeast, KNL-2 in *C. elegans* and plants, Mis18 complex in vertebrates). Through two-hybrid and in vitro biochemical assays, we provide evidence that the *C. elegans* CENP-A N-Tail possesses the latter activity—it interacts directly with the middle region of KNL-2 and this interaction is preserved in a species-specific manner in *C. briggsae,* despite significant primary sequence divergence (especially in the N-Tail sequence). A recent study independently described a CENP-A^HCP-3^ N-tail – KNL-2 interaction that is consistent with what we report here (Prosée et al., 2020). Unfortunately, we have been unable to selectively mutate this interaction and assess the consequences *in vivo*—this will be important future work. An obvious question emerging from our results is whether the N-tail of CENP-A^HCP-3^ also exhibits chaperone activity, analogous to HJURP/Scm3. In preliminary work, we have not observed an interaction between the N-Tail and the histone-fold of CENP-A^HCP-3^ in two-hybrid and in vitro binding assays, which would argue against presence of chaperone activity. However, significant more effort needs to be placed on reconstitutions with purified components to address whether these negative results are indeed due to absence of chaperone activity. RNAi experiments have implicated the *C. elegans* ortholog of the histone-binding WD40 domain chaperone RbAp46/48, LIN-53, in CENP-A chromatin assembly, suggesting that it may work together with the N-tail – KNL-2 interaction described here to assemble centromeric chromatin (Lee et al., 2016).

In conclusion, we provide evidence for an evolutionary variation on CENP-A chromatin assembly in which the N-tail of CENP-A has acquired part of the function of the specialized chaperone/targeting factor HJURP/Scm3 and become essential for loading onto chromatin. This represents a distinct solution from *Drosophila,* which also lacks HJURP/Scm3 but appears to have convergently evolved an HJURP/Scm3-like chaperone called Cal1 (Chen et al., 2014; Erhardt et al., 2008; Medina-Pritchard et al., 2020; Phansalkar et al., 2012). Understanding how different species build CENP-A chromatin at restricted genomic locations should provide insight into the general principles by which the epigenetic state of centromeric chromatin is defined and propagated.

## MATERIALS AND METHODS

### C. elegans strains

*C. elegans* strains (genotypes in *Table S1)* were maintained at 20°C. Engineered GFP::CENP-A^HCP-3^ transgenes were cloned into pCFJ151 and injected into strain EG4322. The KNL-1::mCherry transgene was cloned into pCFJ178 and injected into EG6700 (Frokjaer-Jensen et al., 2008). The amplified *hcp-3* genomic locus was flanked on the 5’ end by 5’-GACGACGCTCCGAATCATTTGGGAG-3’and on the 3’ end by 5’- CTATTTGTCAAATAATAAAGATTCATTACTTGTAAATGAGAACATTTTATTTAA-3’. For the GFP:: CENP-A^HCP-3^ transgenes the GFP sequence was inserted following the start codon and preceded by a GGRAGSGGRAGSGGRAGS linker. All exons were reencoded in *CENP-A^hcp-3^* to allow RNAi-mediated depletion of endogenous CENP-A without affecting the introduced transgene. Single copy insertion was confirmed by PCR. Transgenic strains were crossed into various marker or deletion strains using standard genetic procedures.

### RNA-mediated interference (RNAi)

Double-stranded RNAs were generated using oligos *(Table S2)* to amplify regions from N2 genomic DNA or cDNA. PCR reactions were used as templates for *in vitro* RNA production (Ambion), and the RNA was purified using a MegaClear kit (Ambion). Eluted RNA from the T3 and T7 reactions were mixed together, combined with 3x soaking buffer (32.7 mM Na2HPO4, 16.5 mM KH2PO4, 6.3 mM KCl, 14.1 mM NH4Cl), and annealed (68°C for 10 min., 37°C for 30 min). dsRNA was injected into L3/L4 hermaphrodite worms 38-42 hours prior to imaging. For double depletions dsRNAs were mixed in equal amounts (≥1-3 mg/ml for each RNA).

### Immunoblotting

For immunoblotting a mixed population of worms growing at 20°C on an NGM+OP50 agar plate were collected with M9+0.1% TritonX-100, pelleted, and washed. Worms were vortexed in a mix of 100 μL M9+0.1% TritonX-100, 50 μL 4x sample buffer, and 100 μL glass beads and boiled then vortexed and boiled again. Samples were run on an SDS-PAGE gel, transferred to a PVDF membrane, probed with 1 μg/ml affinity-purified anti-CENP-A^HCP-3^ (rabbit; antigen was CENP-A^HCP-3^(105-183)::6xHis) and detected using an HRP-conjugated secondary antibody (rabbit or mouse; GE Healthcare). For antibody production CENP-A^HCP-3^(105-183)::6xHis was expressed in *E. coli,* purified, and injected into rabbits (Covance). Serum was affinity purified on a HiTrap NHS column to which CENP-A^HCP-3^ (105-183)::6xHis was covalently coupled.

### Yeast Two Hybrid Screens

Yeast two hybrid analysis was performed according to the manufacturer guidelines (Matchmaker; Clontech Laboratories, Inc.). Genes of interest were cloned from wildtype (N2) *C. elegans* or *C. briggsae* cDNA.

### Imaging and Quantification

For all experiments images were acquired using an inverted Zeiss Axio Observer Z1 system with a Yokogawa spinning-disk confocal head (CSU-X1), a 63x 1.4 NA Plan Apochromat objective, and a QuantEM 512SC EMCCD camera (Photometrics). Environmental temperatures during experimental acquisitions averaged 19°C.

For live imaging of one-cell embryos, gravid hermaphrodite adult worms were dissected into M9 buffer, embryos were manually transferred to 2% agarose pads, and overlaid with a coverslip. To monitor chromosome localizations a 5×2 μm z-series was collected every 10-15s in one-cell embryos.

All images and movies were processed, scaled, and analyzed using ImageJ (Fiji), and Photoshop (Adobe). Quantification of CENP-A^HCP-3^ and KNL-1 kinetochore localization during metaphase of one-cell embryos was performed on maximum intensity projections. A rectangle was drawn around the fluorescence signal and average pixel intensity was measured. The rectangle was expanded on all sides by a few pixels and the difference in integrated intensity between the expanded rectangle and the original rectangle was used to define the background intensity per pixel. Integrated fluorescence was then calculated for the original rectangle after background subtraction (Moyle et al., 2014).

### Protein Purification

CENP-A^HCP-3^(1-109)-MBP-6xHis was cloned into pET21a. HCP-3(1-109)-MBP-6xHis and MBP::6xHis were expressed in BL21(DE3). *E. coli* cultures were grown to OD_600_ 0.6-0.8 and induced with 0.1 mM IPTG for 6 hours at 20°C. Induced BL21 (DE3) cells were lysed in Lysis Buffer (20 mM Tris [pH 7.5], 300 mM NaCl, 20 mM imidazole, 8 mM β-mercaptoethanol [BME]) and clarified at 40,000 g for 45 min at 4°C. Ni-NTA agarose (Qiagen) was incubated with clarified lysates for 45 min, washed with Wash Buffer (20 mM Tris [pH 7.5], 300 mM NaCl, 50 mM imidazole, 8 mM BME), and eluted with 20 mM Tris [pH 7.5], 300 mM NaCl, 300 mM imidazole, 8 mM BME. The eluted protein was fractionated using a Superose10 gel filtration column (GE Healthcare). Protein concentrations were determined using a NanoDrop 1000 spectrophotometer (Thermo Scientific).

### In vitro Translation and Binding Assay

KNL-2-MBP fragments were ^[35]^S labeled using TnT^®^ Quick Coupled Transcription/Translation System (Promega). 10 μl of the in vitro translation lysate was incubated for 1hr at 4°C with 50 μg CENP-A^HCP-3^(1-109)-MBP-His (in 20 mM Tris [pH7.5], 300 mM Nacl, 0.05 % NP40, 10 mM Imidazole) in a final volume of 50 μl, mixed with 25 μl of a 1:1 nickel agarose slurry equilibrated with the binding buffer for an additional hour at 4°C. Beads were washed three times with binding buffer, eluted using sample buffer and the elution analyzed by SDS-PAGE and autoradiography.

## ACKNOWLEDGMENTS

This work was supported by an NIH grant (GM074215) to A.D., a German Research Foundation (DFG) grant (GR 3859/1-1) to C.D.G., a NSF Graduate Research Fellowship to J.H., and an EMBO Fellowship (ALTF 251-2012) to A.G-G. A.D., K.O. and A.K.S. received salary and other support from Ludwig Cancer Research.

## SUPPLEMENTARY FIGURE LEGENDS

**Supplementary Figure 1.**
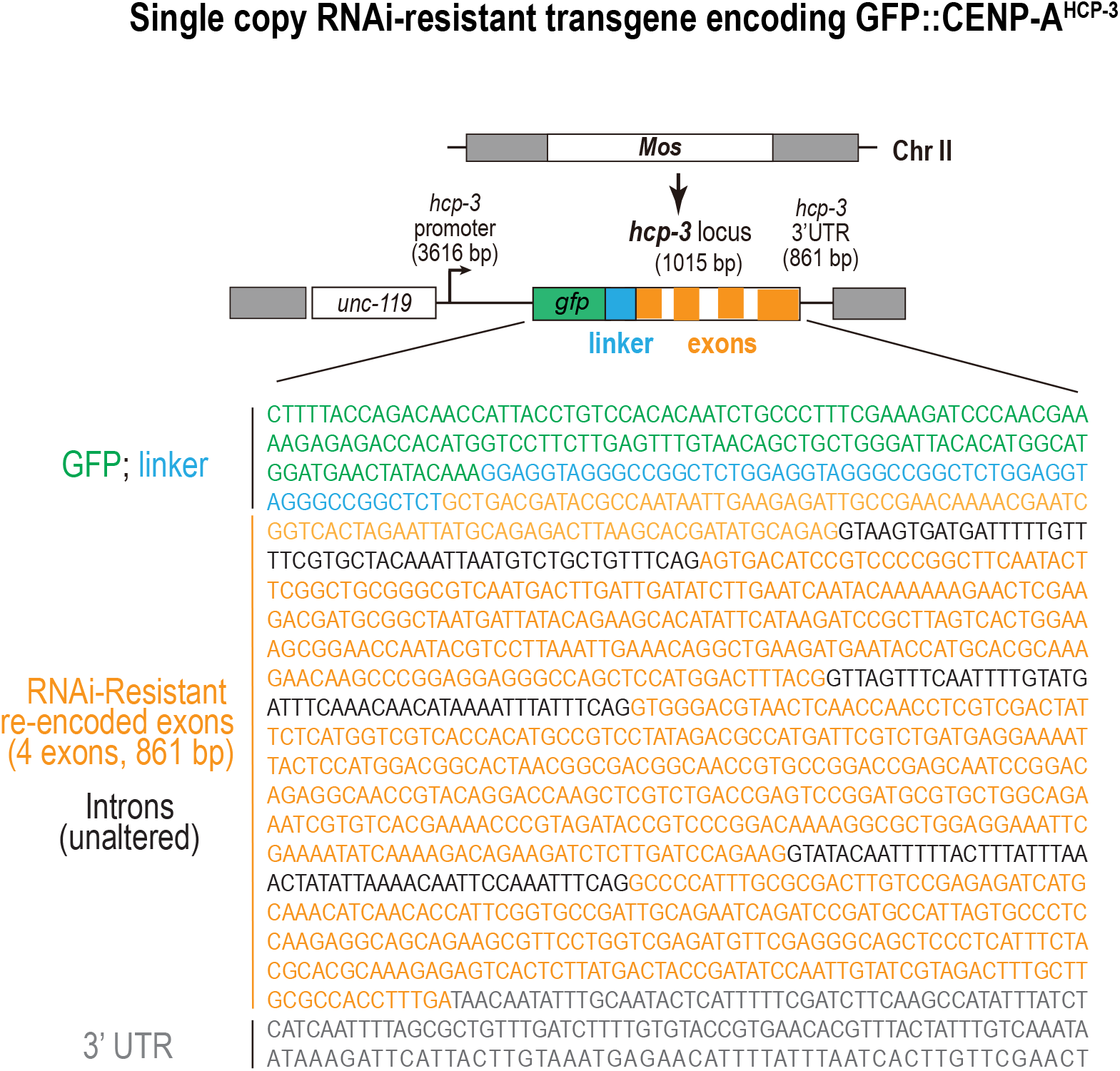
Design of the RNAi-resistant transgene used to replace endogenous CENP-A^HCP-3^. Schematic of the *gfp::hcp-3* RNAi-resistant single copy transgene. All of the exons of *hcp-3* were re-encoded to preserve amino acid sequence but make the nucleotide sequence resistant to a dsRNA generated using the *hcp-3* cDNA. The intron sequences *(indicated in black)* were not altered.

**Supplementary Figure 2.**
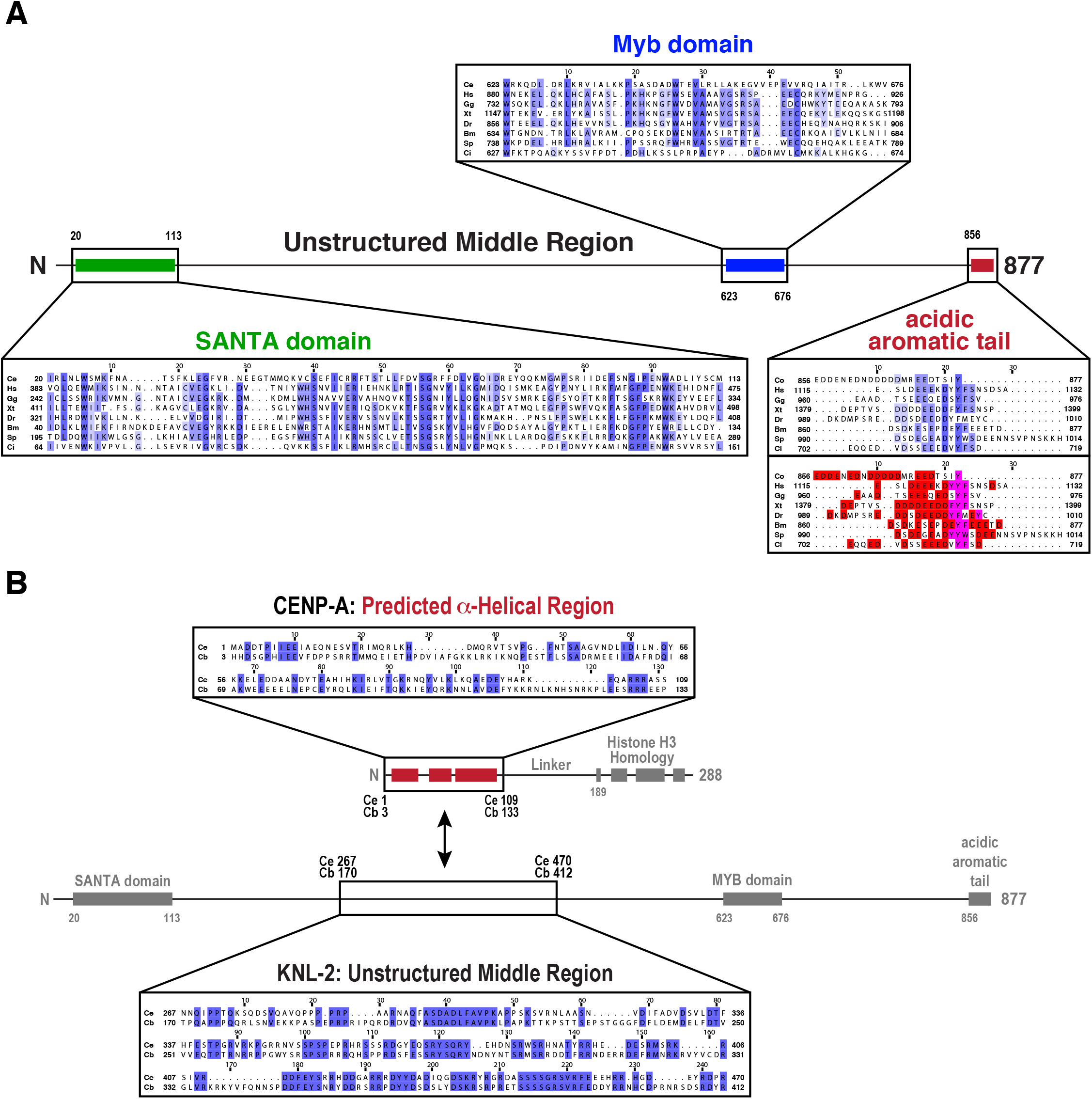
Sequence alignment of KNL-2 protein family domains & *C. elegans/C. briggsae* CENP-A N-tail predicted a-helical regions. **(A)** Alignment of KNL-2/M18BP1 domains from the following species: Ce, *Caenorhabditis elegans;* Hs, *Homo sapiens;* Gg, *Gallus gallus;* Xt, *Xenopus tropicalis;* Dr, *Danio rerio; Bm, Brugia malayi;* Sp, *Strongylocentrotus purpuratus;* Ci, *Ciona intestinalis.* Dark blue lines indicate identical residues and lighter blue lines indicate similar residues among different species. Red and magenta-colored residues in the bottom panel of the C-terminal end highlight acidic and aromatic residues, respectively. Sequences alignments were performed using Clustal Omega. **(B)** Alignment of *C. elegans* and *C. briggsae* predicted N-tail a-helical region and of the interacting middle region from KNL-2.

**Table S1.** *C. elegans* strains used in this study.

| **Strain Number** | **Genotype** |
| --- | --- |
| N2 | Ancestral |
| OD56 | *unc-119(ed3)III; ltIs37 [pAA64; Ppie-1::mCherry::his-58; unc-119 (+)]IV* |
| OD704 | *unc-119(ed3) III; ltSi396[pOD1368; gfp::hcp-3 reencoded; cb unc-119 (+)]II* |
| OD1290 | *unc-119(ed3) III; ltSi409[pOD1369; gfp::hcp-3(Δ109) reencoded; cb unc-119 (+)]II* |
| OD1283 | *unc-119(ed3) III; ltSi406[pOD1372; gfp::hcp-3(Δ184) reencoded]; cb unc-119 (+)]II* |
| OD1271 | *unc-119(ed3) III; ltSi400[pOD1366; knl-1 reencoded::mCherry; cb unc-119 (+)]IV* |
| OD2450 | *unc-119(ed3) III; ltSi396[pOD1368; gfp::hcp-3 reencoded; cb unc-119 (+)]II; ltIs37 [pAA64; Ppie-1::mCherry::his-58; unc-119 (+)]IV* |
| OD1292 | *unc-119(ed3) III; ltSi409[pOD1369; gfp::hcp-3(Δ109) reencoded; cb unc-119 (+)]II; ltIs37 [pAA64; Ppie-1::mCherry::his-58; unc-119 (+)]IV* |
| OD2451 | *unc-119(ed3) III; ltSi396[pOD1368; gfp::hcp-3 reencoded; cb unc-119 (+)]II; ltSi400[pOD1366; knl-1 reencoded::mCherry; cb unc-119 (+)]IV* |
| OD1291 | *unc-119(ed3) III; ltSi409[pOD1369; gfp::hcp-3 reencoded(Mutant 110-288); cb unc-119 (+)]II; ltSi400[pOD1366; knl-1 reencoded::mCherry; cb unc-119 (+)]IV* |
| OD785 | *unc-119(ed3) III; ltSi396[pOD1368; gfp::hcp-3 reencoded; cb unc-119 (+)]II; hcp-3(ok1892) III* |

**Table S2.**
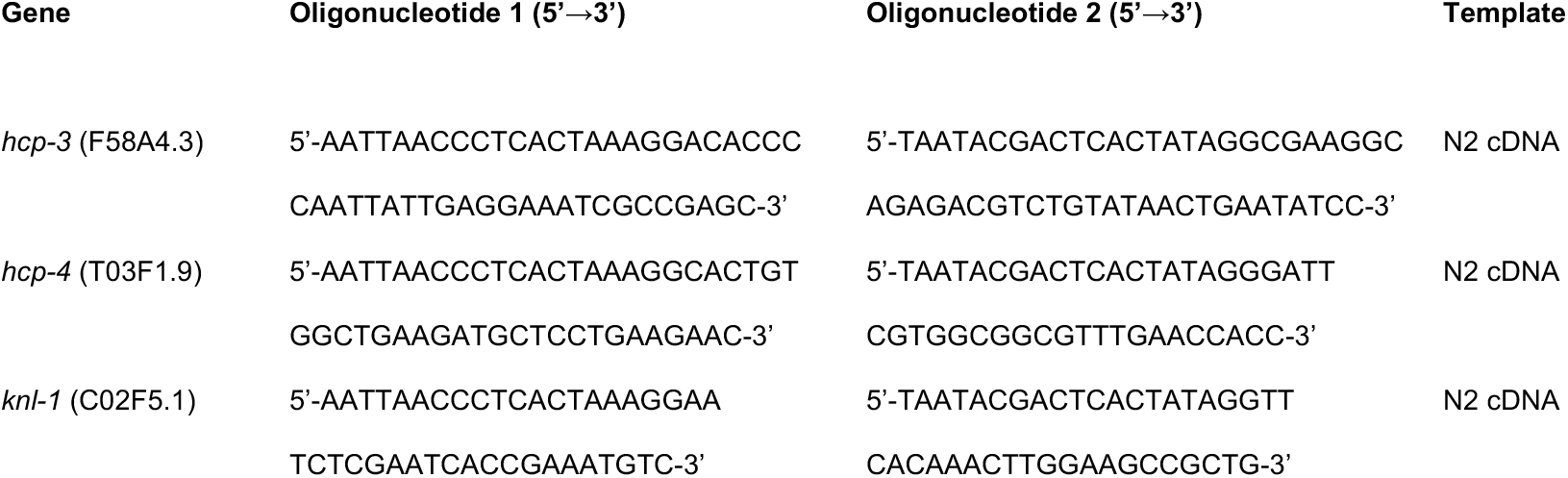
Oligos and templates used for dsRNA production.

## References

Allshire, R.C., and G.H. Karpen. 2008. Epigenetic regulation of centromeric chromatin: old dogs, new tricks? Nat Rev Genet. 9:923–937.

Barnhart, M.C., P.H. Kuich, M.E. Stellfox, J.A. Ward, E.A. Bassett, B.E. Black, and D.R. Foltz. 2011. HJURP is a CENP-A chromatin assembly factor sufficient to form a functional de novo kinetochore. J Cell Biol. 194:229–243.

Black, B.E., D.R. Foltz, S. Chakravarthy, K. Luger, V.L. Woods, Jr., and D.W. Cleveland. 2004. Structural determinants for generating centromeric chromatin. Nature. 430:578–582.

Black, B.E., L.E. Jansen, P.S. Maddox, D.R. Foltz, A.B. Desai, J.V. Shah, and D.W. Cleveland. 2007. Centromere identity maintained by nucleosomes assembled with histone H3 containing the CENP-A targeting domain. Mol Cell. 25:309–322.

Boxem, M., Z. Maliga, N. Klitgord, N. Li, I. Lemmens, M. Mana, L. de Lichtervelde, J.D. Mul, D. van de Peut, M. Devos, N. Simonis, M.A. Yildirim, M. Cokol, H.L. Kao, A.S. de Smet, H. Wang, A.L. Schlaitz, T. Hao, S. Milstein, C. Fan, M. Tipsword, K. Drew, M. Galli, K. Rhrissorrakrai, D. Drechsel, D. Koller, F.P. Roth, L.M. Iakoucheva, A.K. Dunker, R. Bonneau, K.C. Gunsalus, D.E. Hill, F. Piano, J. Tavernier, S. van den Heuvel, A.A. Hyman, and M. Vidal. 2008. A protein domain-based interactome network for C. elegans early embryogenesis. Cell. 134:534–545.

Chen, C.C., M.L. Dechassa, E. Bettini, M.B. Ledoux, C. Belisario, P. Heun, K. Luger, and B.G. Mellone. 2014. CAL1 is the Drosophila CENP-A assembly factor. J Cell Biol. 204:313–329.

Chen, Y., R.E. Baker, K.C. Keith, K. Harris, S. Stoler, and M. Fitzgerald-Hayes. 2000. The N terminus of the centromere H3-like protein Cse4p performs an essential function distinct from that of the histone fold domain. Mol Cell Biol. 20:7037–7048.

Cho, U.S., and S.C. Harrison. 2011. Recognition of the centromere-specific histone Cse4 by the chaperone Scm3. Proc Natl Acad Sci U S A. 108:9367–9371.

Desai, A., S. Rybina, T. Muller-Reichert, A. Shevchenko, A. Shevchenko, A. Hyman, and K. Oegema. 2003. KNL-1 directs assembly of the microtubule-binding interface of the kinetochore in C. elegans. Genes Dev. 17:2421–2435.

Dunleavy, E.M., D. Roche, H. Tagami, N. Lacoste, D. Ray-Gallet, Y. Nakamura, Y. Daigo, Y. Nakatani, and G. Almouzni-Pettinotti. 2009. HJURP is a cell-cycle-dependent maintenance and deposition factor of CENP-A at centromeres. Cell. 137:485–497.

Erhardt, S., B.G. Mellone, C.M. Betts, W. Zhang, G.H. Karpen, and A.F. Straight. 2008. Genomewide analysis reveals a cell cycle-dependent mechanism controlling centromere propagation. J Cell Biol. 183:805–818.

Espeut, J., D.K. Cheerambathur, L. Krenning, K. Oegema, and A. Desai. 2012. Microtubule binding by KNL-1 contributes to spindle checkpoint silencing at the kinetochore. J Cell Biol. 196:469–482.

Fachinetti, D., H.D. Folco, Y. Nechemia-Arbely, L.P. Valente, K. Nguyen, A.J. Wong, Q. Zhu, A.J. Holland, A. Desai, L.E. Jansen, and D.W. Cleveland. 2013. A two-step mechanism for epigenetic specification of centromere identity and function. Nat Cell Biol. 15:1056–1066.

Folco, H.D., C.S. Campbell, K.M. May, C.A. Espinoza, K. Oegema, K.G. Hardwick, S.I.S. Grewal, and A. Desai. 2015. The CENP-A N-tail confers epigenetic stability to centromeres via the CENP-T branch of the CCAN in fission yeast. Curr Biol. 25:348–356.

Foltz, D.R., L.E. Jansen, A.O. Bailey, J.R. Yates, 3rd, E.A. Bassett, S. Wood, B.E. Black, and D.W. Cleveland. 2009. Centromere-specific assembly of CENP-a nucleosomes is mediated by HJURP. Cell. 137:472–484.

French, B.T., and A.F. Straight. 2019. CDK phosphorylation of Xenopus laevis M18BP1 promotes its metaphase centromere localization. EMBO J. 38.

French, B.T., F.G. Westhorpe, C. Limouse, and A.F. Straight. 2017. Xenopus laevis M18BP1 Directly Binds Existing CENP-A Nucleosomes to Promote Centromeric Chromatin Assembly. Dev Cell. 42:190–199 e110.

Frokjaer-Jensen, C., M.W. Davis, C.E. Hopkins, B.J. Newman, J.M. Thummel, S.P. Olesen, M. Grunnet, and E.M. Jorgensen. 2008. Single-copy insertion of transgenes in Caenorhabditis elegans. Nat Genet. 40:1375–1383.

Fujita, Y., T. Hayashi, T. Kiyomitsu, Y. Toyoda, A. Kokubu, C. Obuse, and M. Yanagida. 2007. Priming of centromere for CENP-A recruitment by human hMis18alpha, hMis18beta, and M18BP1. Dev Cell. 12:17–30.

Gassmann, R., A. Rechtsteiner, K.W. Yuen, A. Muroyama, T. Egelhofer, L. Gaydos, F. Barron, P. Maddox, A. Essex, J. Monen, S. Ercan, J.D. Lieb, K. Oegema, S. Strome, and A. Desai. 2012. An inverse relationship to germline transcription defines centromeric chromatin in C. elegans. Nature. 484:534–537.

Hayashi, T., Y. Fujita, O. Iwasaki, Y. Adachi, K. Takahashi, and M. Yanagida. 2004. Mis16 and Mis18 are required for CENP-A loading and histone deacetylation at centromeres. Cell. 118:715–729.

Hori, T., W.H. Shang, M. Hara, M. Ariyoshi, Y. Arimura, R. Fujita, H. Kurumizaka, and T. Fukagawa. 2017. Association of M18BP1/KNL2 with CENP-A Nucleosome Is Essential for Centromere Formation in Non-mammalian Vertebrates. Dev Cell. 42:181–189 e183.

Hu, H., Y. Liu, M. Wang, J. Fang, H. Huang, N. Yang, Y. Li, J. Wang, X. Yao, Y. Shi, G. Li, and R.M. Xu. 2011. Structure of a CENP-A-histone H4 heterodimer in complex with chaperone HJURP. Genes Dev. 25:901–906.

Jansen, L.E., B.E. Black, D.R. Foltz, and D.W. Cleveland. 2007. Propagation of centromeric chromatin requires exit from mitosis. J Cell Biol. 176:795–805.

Kato, H., J. Jiang, B.R. Zhou, M. Rozendaal, H. Feng, R. Ghirlando, T.S. Xiao, A.F. Straight, and Y. Bai. 2013. A conserved mechanism for centromeric nucleosome recognition by centromere protein CENP-C. Science. 340:1110–1113.

Kixmoeller, K., P.K. Allu, and B.E. Black. 2020. The centromere comes into focus: from CENP-A nucleosomes to kinetochore connections with the spindle. Open Biol. 10:200051.

Lee, B.C., Z. Lin, and K.W. Yuen. 2016. RbAp46/48(LIN-53) Is Required for Holocentromere Assembly in Caenorhabditis elegans. Cell Rep. 14:1819–1828.

Lermontova, I., M. Kuhlmann, S. Friedel, T. Rutten, S. Heckmann, M. Sandmann, D. Demidov, V. Schubert, and I. Schubert. 2013. Arabidopsis kinetochore null2 is an upstream component for centromeric histone H3 variant cenH3 deposition at centromeres. Plant Cell. 25:3389–3404.

Maddox, P.S., F. Hyndman, J. Monen, K. Oegema, and A. Desai. 2007. Functional genomics identifies a Myb domain-containing protein family required for assembly of CENP-A chromatin. J Cell Biol. 176:757–763.

Maddox, P.S., N. Portier, A. Desai, and K. Oegema. 2006. Molecular analysis of mitotic chromosome condensation using a quantitative time-resolved fluorescence microscopy assay. Proc Natl Acad Sci U S A. 103:15097–15102.

McKinley, K.L., and I.M. Cheeseman. 2016. The molecular basis for centromere identity and function. Nat Rev Mol Cell Biol. 17:16–29.

Medina-Pritchard, B., V. Lazou, J. Zou, O. Byron, M.A. Abad, J. Rappsilber, P. Heun, and A.A. Jeyaprakash. 2020. Structural basis for centromere maintenance by Drosophila CENP-A chaperone CAL1. EMBO J. 39:e103234.

Mellone, B.G., K.J. Grive, V. Shteyn, S.R. Bowers, I. Oderberg, and G.H. Karpen. 2011. Assembly of Drosophila centromeric chromatin proteins during mitosis. PLoS Genet. 7:e1002068.

Melters, D.P., L.V. Paliulis, I.F. Korf, and S.W. Chan. 2012. Holocentric chromosomes: convergent evolution, meiotic adaptations, and genomic analysis. Chromosome Res. 20:579–593.

Mitra, S., B. Srinivasan, and L.E.T. Jansen. 2020. Stable inheritance of CENP-A chromatin: Inner strength versus dynamic control. J Cell Biol. 219.

Moree, B., C.B. Meyer, C.J. Fuller, and A.F. Straight. 2011. CENP-C recruits M18BP1 to centromeres to promote CENP-A chromatin assembly. J Cell Biol. 194:855–871.

Moyle, M.W., T. Kim, N. Hattersley, J. Espeut, D.K. Cheerambathur, K. Oegema, and A. Desai. 2014. A Bub1-Mad1 interaction targets the Mad1-Mad2 complex to unattached kinetochores to initiate the spindle checkpoint. J Cell Biol. 204:647–657.

Musacchio, A., and A. Desai. 2017. A Molecular View of Kinetochore Assembly and Function. Biology (Basel). 6.

Oegema, K., A. Desai, S. Rybina, M. Kirkham, and A.A. Hyman. 2001. Functional analysis of kinetochore assembly in Caenorhabditis elegans. J Cell Biol. 153:1209–1226.

Ohzeki, J., J.H. Bergmann, N. Kouprina, V.N. Noskov, M. Nakano, H. Kimura, W.C. Earnshaw, V. Larionov, and H. Masumoto. 2012. Breaking the HAC Barrier: histone H3K9 acetyl/methyl balance regulates CENP-A assembly. EMBO J. 31:2391–2402.

Pan, D., K. Walstein, A. Take, D. Bier, N. Kaiser, and A. Musacchio. 2019. Mechanism of centromere recruitment of the CENP-A chaperone HJURP and its implications for centromere licensing. Nat Commun. 10:4046.

Phansalkar, R., P. Lapierre, and B.G. Mellone. 2012. Evolutionary insights into the role of the essential centromere protein CAL1 in Drosophila. Chromosome Res. 20:493–504.

Pidoux, A.L., E.S. Choi, J.K. Abbott, X. Liu, A. Kagansky, A.G. Castillo, G.L. Hamilton, W. Richardson, J. Rappsilber, X. He, and R.C. Allshire. 2009. Fission yeast Scm3: A CENP-A receptor required for integrity of subkinetochore chromatin. Mol Cell. 33:299–311.

Prosée, R.F., J.M. Wenda, C. Gabus, K. Delaney, F. Schwager, M. Gotta, and F.A. Steiner. 2020. Trans-generational inheritance of centromere identity requires the CENP-A N-terminal tail in the *C. elegans* maternal germ line. bioRxiv.

Ravi, M., P.N. Kwong, R.M. Menorca, J.T. Valencia, J.S. Ramahi, J.L. Stewart, R.K. Tran, V. Sundaresan, L. Comai, and S.W. Chan. 2010. The rapidly evolving centromere-specific histone has stringent functional requirements in Arabidopsis thaliana. Genetics. 186:461–471.

Sanchez-Pulido, L., A.L. Pidoux, C.P. Ponting, and R.C. Allshire. 2009. Common ancestry of the CENP-A chaperones Scm3 and HJURP. Cell. 137:1173–1174.

Shuaib, M., K. Ouararhni, S. Dimitrov, and A. Hamiche. 2010. HJURP binds CENP-A via a highly conserved N-terminal domain and mediates its deposition at centromeres. Proc Natl Acad Sci U S A. 107:1349–1354.

Steiner, F.A., and S. Henikoff. 2014. Holocentromeres are dispersed point centromeres localized at transcription factor hotspots. Elife. 3:e02025.

Wang, J., X. Liu, Z. Dou, L. Chen, H. Jiang, C. Fu, G. Fu, D. Liu, J. Zhang, T. Zhu, J. Fang, J. Zang, J. Cheng, M. Teng, X. Ding, and X. Yao. 2014. Mitotic regulator Mis18beta interacts with and specifies the centromeric assembly of molecular chaperone holliday junction recognition protein (HJURP). J Biol Chem. 289:8326–8336.

Williams, J.S., T. Hayashi, M. Yanagida, and P. Russell. 2009. Fission yeast Scm3 mediates stable assembly of Cnp1/CENP-A into centromeric chromatin. Mol Cell. 33:287–298.

Zhang, D., C.J. Martyniuk, and V.L. Trudeau. 2006. SANTA domain: a novel conserved protein module in Eukaryota with potential involvement in chromatin regulation. Bioinformatics. 22:2459–2462.

